# Calculating sample size for identifying cell subpopulation in single-cell RNA-seq experiments

**DOI:** 10.1101/706481

**Authors:** Kyung In Kim, Ahrim Youn, Mohan Bolisetty, A. Karolina Palucka, Joshy George

## Abstract

Single-cell RNA sequencing (scRNA-seq) is a rapidly developing technology for studying gene expression at the individual cell level and is often used to identify subpopulations of cells. Although the use of scRNA-seq is steadily increasing in basic and translational research, there is currently no statistical model for calculating the optimal number of cells for use in experiments that seek to identify cell subpopulations. Here, we have developed a statistical method ***ncells*** for calculating the number of cells required to detect a rare subpopulation in a homogeneous cell population for the given type I and II error. ***ncells*** defines power as the probability of separation of subpopulations which is calculated from three user-defined parameters: the proportion of rare subpopulation, proportion of up-regulated marker genes of the subpopulation, and levels of differential expression of the marker genes. We applied ***ncells*** to the scRNA-seq data on dendritic cells and monocytes isolated from healthy blood donor to show its efficacy in calculating the optimal number of cells in identifying a novel subpopulation.

## 1. Itroduction

Single-cell RNA sequencing (scRNA-seq) technology (Tang *and others*, 2009; Ramsköld *and others*, 2012) has been rapidly applied to many biomedical research areas such as cancer (Patel *and others*, 2014), neurobiology (Tasic *and others*, 2016) and immunology (Villani *and others*, 2017). One of the main applications is classifying single cells of a given tissue into specific cell subpopulations based on the genes they express. Single cell classification has been useful in identifying a new cell subtypes to refine existing classification (Villani *and others*, 2017). While such classification has been successfully accomplished through the development of numerous computational methods in recent years (Bacher and Kendziorski, 2016), no statistical method has been developed for calculating the optimal number of cells that would allow identifying a new cell subtype.

There are mainly two challenges in calculating the number of cells required for identifying a new cell type. First, identifying a novel subpopulation from a known cell type population relies on unsupervised clustering methods such as principal component analysis (PCA), t-distributed stochastic neighbor embedding (t-SNE) (Maaten and Hinton, 2008) and UMAP (McInnes and Healy, 2018) while traditional sample size calculation is performed in the supervised hypothesis-testing framework (Hart *and others*, 2013).

Second, the statistical properties of scRNA-seq data are markedly different from those of bulk RNA-seq data. In particular, the number of genes with a zero read count is much higher in datasets for scRNA-seq than in bulk RNA-seq due to technical and/or biological reasons (Kharchenko *and others*, 2014). These dropout events need to be modeled properly to calculate an appropriate cell sample size.

Here we propose a statistical model of scRNA-seq data for calculating appropriate number of cells for discovering a novel, rare cell type in a seemingly homogeneous population, which takes into account the characteristics of scRNA-seq data such as dropout events. Using this model, we estimated the population mean difference by projecting gene expressions onto a separating direction and used a hypothesis-testing framework to determine the separation of the two popuations with relevant type I and type II error. We implemented our methods in an R package called ***ncells*** (https://github.com/TheJacksonLaboratory/ncells). In the Methods, we explain the details of our model, and in the Results, we applied ***ncells*** to the simulated scRNA-seq data and the scRNA-seq data on dendritic cells (DCs) and monocytes isolated from healthy blood donor (Villani *and others*, 2017) to show its efficacy in calculating the optimal number of cells in identifying a novel subpopulation. We also implemented a web-based pre-calculated version of ***ncells*** for selected input parameters that is available at https://ncells.shinyapps.io/ncells.

## 2. Methods

***ncells*** aim to estimate the number of cells required to distinguish a subpopulation from the population of cells that consist of two subpopulations Π_0_ and Π_1_. The subpopulation Π_1_ is separated from Π_0_ using the marker genes which we define as upregulated genes in the subpopulation Π_1_. ***ncells*** use three key parameters that distinguish a subpopulation within a larger cell population: 1. the proportion of novel subpopulation Π_1_; 2. the number of marker genes up-regulated in the subpopulation; 3. strength of marker gene expression (e.g. > 2-fold change increase in expression level). We first model gene expression without dropout event using a hierarchical normal distribution and then incorporate dropout events.

### 2.1 Modeling gene expression level in the absence of dropout event

We assume that log-transformed normalized read count values (e.g. log(*TPM* +1)) are normally distributed (Shalek *and others*, 2014; Law *and others*, 2014). Let *Y_ij_* be the variance-standardized, log-transformed count data of *i*th gene and *j*th cell. We then model expression level of each gene in a hierarchical manner.

For the non-marker gene *i* in cell *j*:

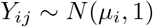

For the marker gene *i*:

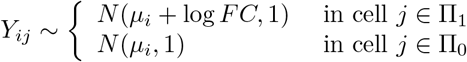

where *FC* is the common fixed fold change of the marker genes. We assume each mean gene expression is distributed as normal with mean *μ* and variance *σ*^2^ as

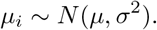

### 2.2 Modeling dropout events

In most scRNA-seq data, high proportions of zero read counts are observed due to biological or technical reasons. The rate of dropout events depends on the sequencing depth and thus varies greatly across different single-cell sequencing platforms. For example, scRNA-seq data obtained using Fludigm C1 is known to have lower dropout rate on average than droplet based platforms such as inDrop or 10X Genomics (Li and Li, 2018). It is also presumed that the dropout event depends on the actual expression level (Marinov *and others*, 2014). Thus we model the probability of dropout event as being inversely proportional to actual expression values with a fixed rate α that takes into account platform-specific dropout rate:

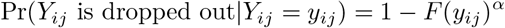

Here, *F* is the marginal cumulative distribution function of *Y_ij_* obtained from the model in Section 2.1 as

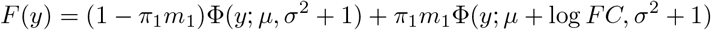

where Φ(*y*; *a, b*) is the normal cumulative distribution with mean *a* and variance *b*. Therefore, dropout event depends both on the expression level and the parameter α that controls overall dropout rate. The overall dropout rate of the data is calculated as:

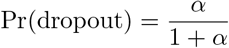

For example, if the overall dropout rate is given as 70% a priori, *α* is set to 7/3.

### 2.3 Detecting a subpopulation

In our model without the dropout event, the difference between the two populations lies in the direction of the first principal component (*PC*1) (see Supplementary Information). With dropout event, *PC*1 still retains the population difference when the mean difference is large (See Supplementary Information for the relation of *PC*1 and the direction between the two population mean vector). Thus we can assess the separation of two populations by measuring the distance between the two sample means projected onto the sample *PC*1 and we define two clusters from the two populations as being separated when

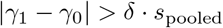

where *γ*_1_, *γ*_0_, *s*_pooled_ are two sample means and the pooled standard deviation of projected values onto *PC*1. Positive *δ* determines the degree of separation between two means *γ*_1_ and *γ*_0_. Figure 1 illustrates an example of two subpopulations separated by *PC*1. We set the value of *δ* so that the type I error Pr(|*γ*_1_ − *γ*_0_| > *δ* · *s*_pooled_|*FC* = 1) equals to the prespecified significance level, for example 0.05. For the more detailed discussion of *PC*1 in separating two populations, see the Supplementary Information.

**Fig. 1.**
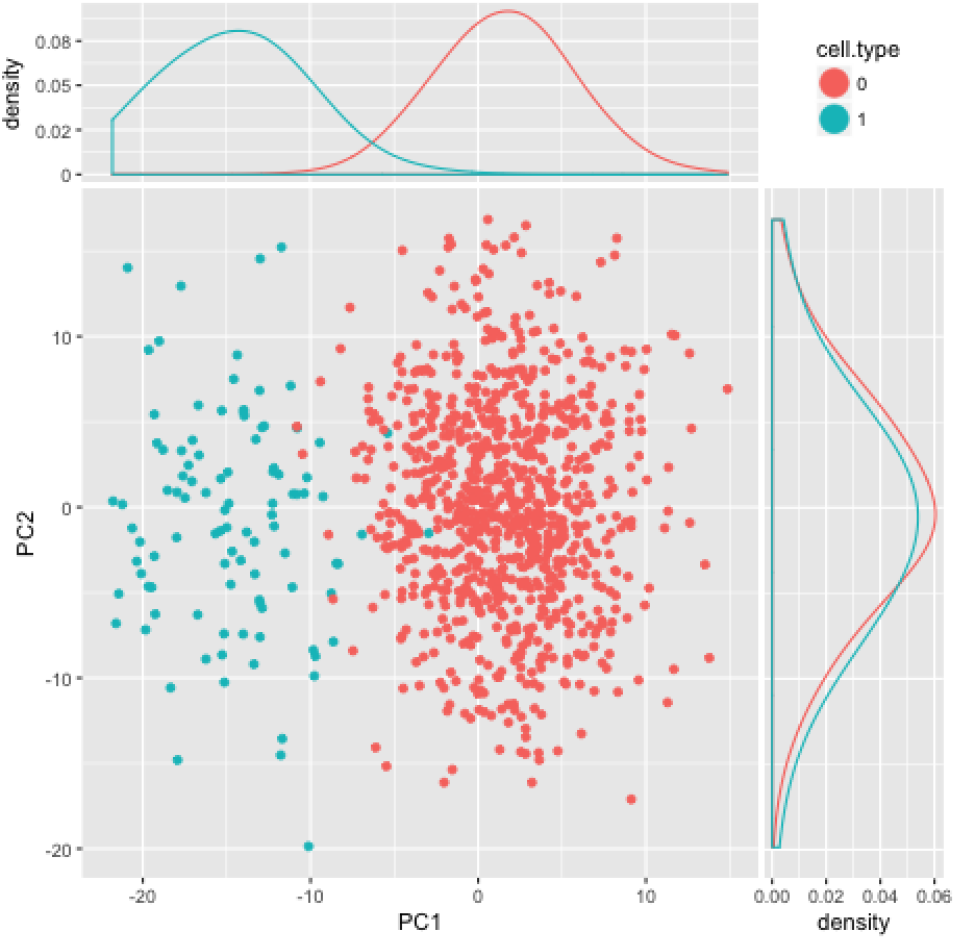
The first two principal component scores of gene expression values of 1000 cells. The first principal component (*PC*1) separates two populations of 900 (orange) and 100 (green) cells while the direction of the second principal component (*PC*2) does not show any separation. Common 2-fold change was applied to 100 marker genes.

### 2.4 Power calculation

Lastly, we estimate the probability of separation of the two populations through simulation, which serves as a power in conventional sample size calculation. The parameters required in the simulation include the mean *μ* and the standard deviation *σ* of the mean expression values, total number of cells sequenced (*n*), the proportion of cells in the subpopulation Π_1_ (*π*_1_), the total number of genes (*p*), the proportion of marker genes (*m*_1_), the common fold change of the marker genes (*FC*), overall dropout rate (*d*) and significance level *α*. For the given parameters, we iterate simulations *B* times and estimate the power (probability of separation) by the frequency of separation of two sample populations out of *B* simulations:

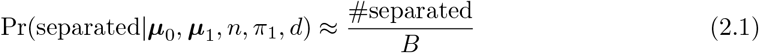

where ***μ***_0_ and ***μ***_1_ are the mean expression vectors of Π_0_ and Π_1_ generated from parameters *μ, σ, m*_1_.

## 3. Results

### 3.1 Simulation study

We performed simulations to assess statistical power to detect new subpopulation within a larger population of cells obtained from scRNA-seq experiment. We randomly selected the first *p* · *m*_1_ genes among *p* genes as the marker genes in the simulation. We simulate gene expression values using our model described in the Methods section with the following parameters.

- Mean *μ* of the mean expression values across genes: 1
- Standard deviation *σ* of the mean expression values across genes: 1
- Total number of cells (*n*): 100, 200,…, 1000
- Proportion of cells in the subpopulation Π_1_ (*π*_1_): 5%, 10%
- Total number of genes (*p*): 20000
- Proportion of marker genes (*m*_1_): 0.1%, 0.2%, 0.4%, 0.8%, 1.6%
- Fold change in marker genes (*FC*): 1.5, 2, 4, 8
- Overall dropout rate (*d*): 60%, 70%, 80%, 90%, 95%

Figure 2 compares the effect of overall dropout rates when the percentage of marker gene is 0.8% with 2 fold change and the percentage of subpopulation = 10%, and *μ* = *σ* = 1. It shows the general pattern of sample size and power: a larger sample size leads to a higher power (higher probability of finding the novel cells). Also as expected, it shows that a higher dropout rate requires more cells to achieve the same level of power. In the web-based ***ncells*** page (http://ncells.shinyapps.io/ncells/), it is shown that power increases by selecting higher marker gene proportions or by increasing fold change. From Figure 2, we can suggest that at least 300 cells would need to be sequenced to identify a subpopulation with 80% power, given 70% dropout rate. For additional results, see the web-based ***ncells*** page.

**Fig. 2.**
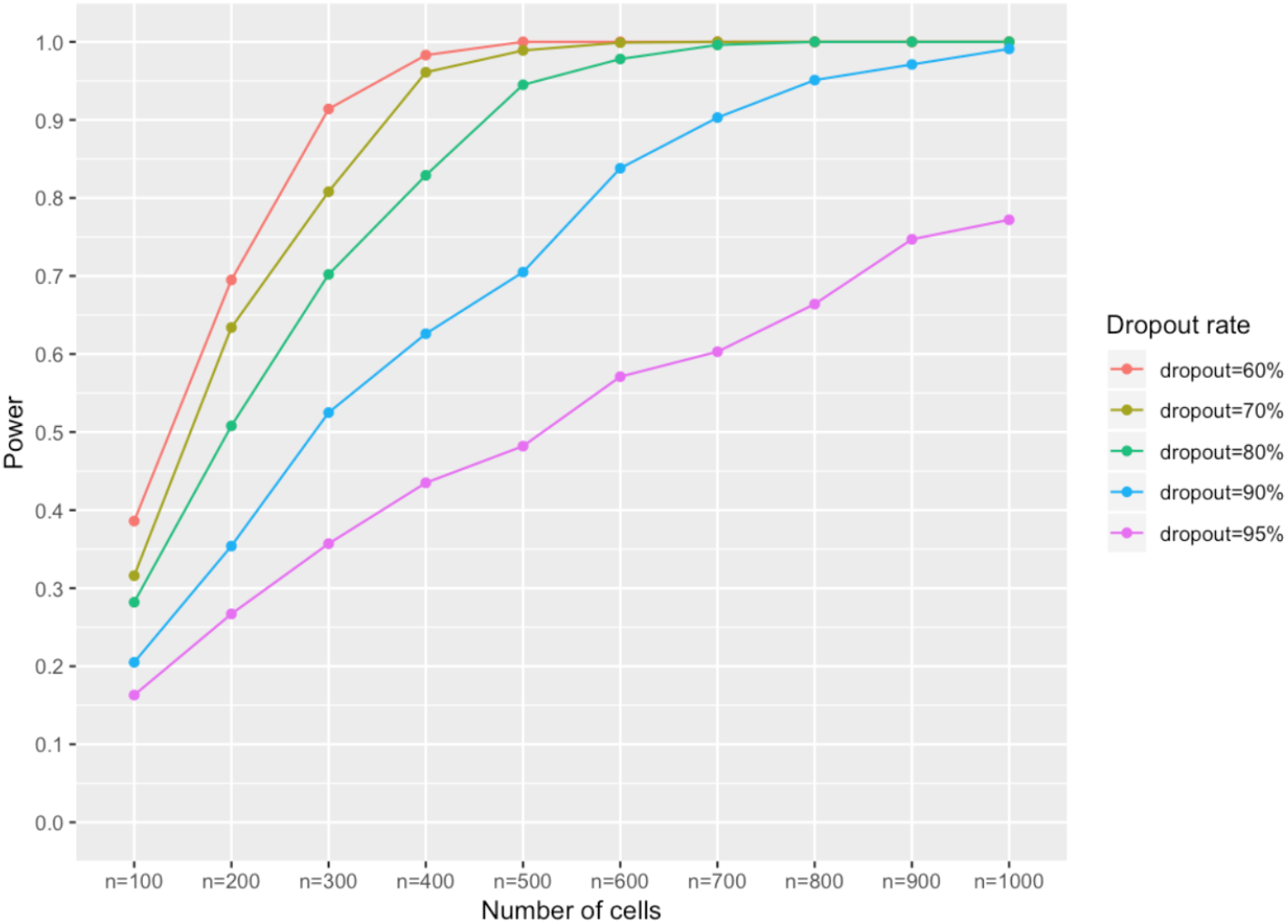
Effect of dropout rate on the relationship between the power and cell sample size. Simulation parameters are: the proportion of marker genes *m*_1_ = 0.8%, the proportion of novel subpopulation *π*_1_ = 10%, 2-fold change in the marker gene expression, mean and variance *μ* = 1, *σ* = 1 for the distribution of mean gene expressions, and total number of gene *p* = 20000.

**Fig. 3.**
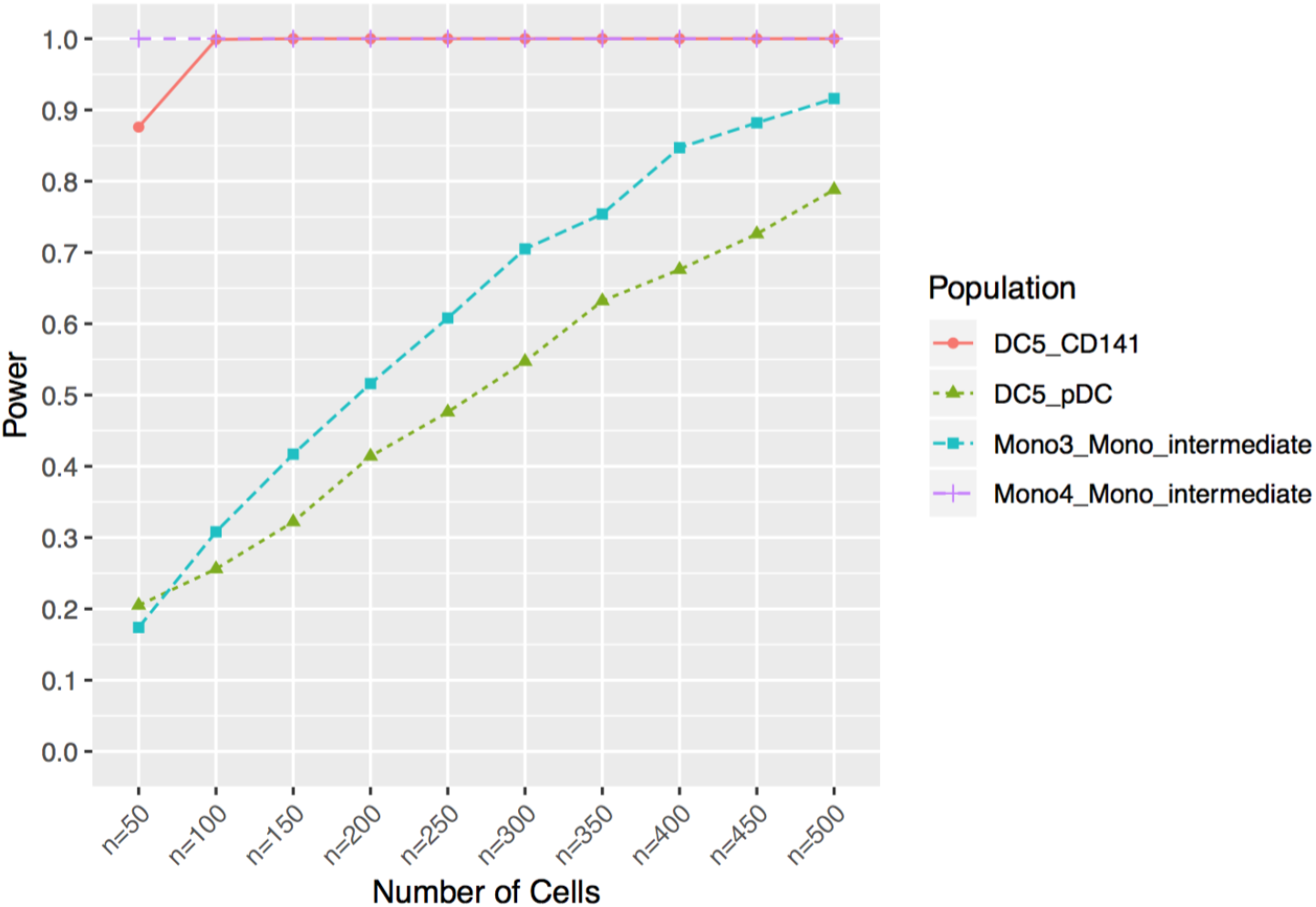
Power and cell sample size for the parameters in Table 1 by ***ncells***. The DC5 cluster in *CD*141 cells and Mono4 cluster in Mono intermediate cells were shown to have sufficiently high power (> 95% for *n* = 188, *n* = 107, respectively). DC5 cluster in pDC cells and Mono3 cluster in Mono intermediate cells were shown to have low power (< 45% for *n* = 182 and ~ 30% for *n* = 107, respectively).

### 3.2 Real data analysis

We applied ***ncells*** to the dataset from Villani *and others* (2017) who performed single-cell RNA sequencing on dendritic cells (*DC*s) and monocytes isolated from a healthy blood donor. Previous studies classified human blood *DC*s into one plasmacytoid *DC* (*pDC*) and two *cDC* populations (*CD*1*C* and *CD*141). By using unsupervised analysis of single cell expression levels, they clustered 742 *DC*s derived from a single donor into 6 clusters, *DC*1, ⋯, *DC*6, one of which is a newly discovered *DC* subpopulation (*DC*5). The *DC*5 population is mixed with the previously classified *CD*141 and *pDC* populations and we applied ***ncells*** to test if *DC*5 can be distinguished within these two populations. They also clustered 339 monocytes into four subpopulations to identify two new monocyte clusters (*Mono*3 and *Mono*4), which belong to the “intermediate” monocyte population gated from *CD*14^++^*CD*16^+^ (hereafter *Mono intermediate* population). Using the parameters estimated from the data obtained from GEO (GSE94820), we estimated power and appropriate number of cells to detect *DC*5 cluster in *CD*141 and *pDC* population and *Mono*3/*Mono*4 cluster in *Mono intermediate* population. The detailed information about the dataset and estimated parameters are in Table 1 and Supplementary Information.

**Table 1.**
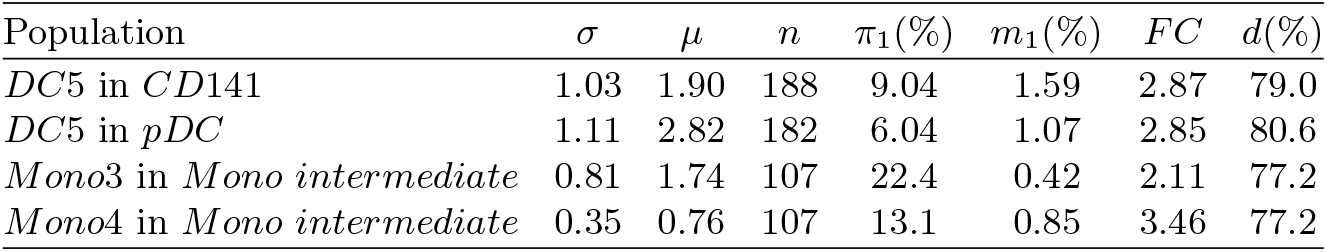
Parameters used to calculate power for detection of the four clusters in Villani and others (2017). edgeR (Robinson and others, 2010) was used to detect differentially expressed genes with FDR 0.05 and positive fold change in the subpopulation. Those differentially expressed genes were used as marker genes. Total number of genes (p) was 26593, estimated mean (μ) and standard deviation (σ) of mean gene expressions, the proportion of cells in the subpopulation (π_1_), the proportion of marker genes (m_1_), common fold change in the marker gene (FC), and overall dropout rate (d) were obtained from the dataset (See Supplementary Information).

We found that two out of the four clusters (*DC*5 in *CD*141, *Mono*4 in *Mono intermediate*) have sufficient powers (> 95%) with total *n* = 188 and *n* = 107 cells, respectively while the other two clusters, *DC*5 in *pDC* and *Mono*3 in *Mono intermediate* didn’t show sufficient power for total *n* = 182 and *n* = 107 cells. Relatively high variance and small proportion of subpopulation (*σ* = 2.82, *π*_1_ = 6.04) may explain the low power of *DC*5 cluster in *pDC* population. For *Mono*3 cluster, the proportion of marker genes (*m*_1_ = 0.42) may explain the low power. Detailed code and information is available in Supplementary information.

## 4. Discussion

We developed a statistical model that incorporates key parameters of scRNA-seq data such as high dropout rate to estimate the number of cells required to identify a cell subpopulation within a seemingly homogeneous population. Within our model, we defined the separation of two populations with the relevant type I and II errors of the hypothesis-testing setup and calculated the optimal number of cells required to attain sufficient power. Our method does not evaluate any particular clustering method so it can be used as a general guide to design scRNA-seq experiment. We implemented the procedure in R package ***ncells*** and also web-based tool for some pre-selected parameters.

We modeled our dropout mechanism with cumulative marginal distribution of gene expression. The key idea was to take into account the protocol-dependent overall dropout rate and to have the dropout event depend only on the actual gene expression level. The overall dropout rate is one crucial factor to determine the dropout effect in sample size calculation. However, there can be alternate approaches (such as logistic regression modeling in Shalek *and others*, 2014) and it would be useful to compare the effect of different dropout mechanisms in the sample size calculation.

We modeled gene expression levels using normal distribution with unit variance for gene exression distribution. Negative binomial model (Robinson *and others*, 2010) or zero-inflated negative binomial model (Van De Wiel *and others*, 2013) have been popular choice for modeling expression values of RNA-seq data. However, those models generally require more parameters to be specified than our model and since simulating high-dimensional gene expression data for many different conditions is computationally demanding task, it is important to reduce the number of parameters while keeping essential properties of scRNA-seq experiment. Further research is needed on more complex models but with computational feasibility.

For very high dropout rate (e.g. > 97%), simulation based sample size calculation method becomes computationally demanding as it requires large number of samples. Recently dropseq based single cell RNA-seq technology (e.g. 10x Genomics) has been generating million cells in an experiment (Zheng *and others*, 2017). In this case, we can consider populaton level inference based on the asymptotic property of our model and estimate the optimal number of cells (See the discussion of Supplementary Information).

***ncells*** is the first statistical method developed for calculating the number of cells required to detect a rare subpopulation in a homogeneous cell population for the given type I and II error. ***ncells*** will be an essential tool in designing single cell RNA sequencing experiment to study cell type classification.

## 5. Software

Software in the form of R code, together with a sample input data set and complete documentation is available on request from the corresponding author.

## Acknowledgments

*Conflict of Interest*: None declared.

## 6. Supplementary Material

Without dropout event, the first principal component direction (*PC*1) is same as the direction that separates the two populations of interest. However, with dropout event when the difference between the two populations is small (e.g. small fold change, small number of marker genes etc), *PC*1 may not represent the population separating direction. Instead it is likely that the *PC*1 indicates the gene having the largest variance among all genes. Thus we investigate the condition for which *PC*1 is the population separating direction. Also we provide relavance of *PC*1 to identify subpopulations for varying parameters in our simulation studies.

### *PC*1 in the absence of dropout event

As discussed in the Method section, we consider two populations Π_0_ and Π_1_ with proportions *π*_0_ and *π*_1_, respectively (*π*_0_ + *π*_1_ = 1). Let *M*_0_ and *M*_1_ be the sets of non-marker and marker genes with proportions *m*_0_, *m*_1_ in total *p* genes (*m*_0_ + *m*_1_ = 1). The gene expression of *i*th gene in *j*th cell was defined as

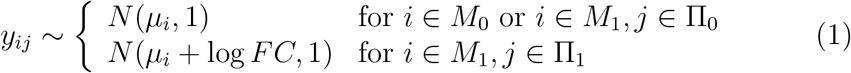

where *μ_i_* is the mean expression of *i*th gene and *FC* is the common fold change for the upregulated genes in *M*_1_. If we let ***Y*** be *p*-vector of gene expressions and ***μ***_0_, ***μ***_1_ be the mean expression vectors of Π_0_, Π_1_, respectively, then the covariance matrix of ***Y*** is derived as:

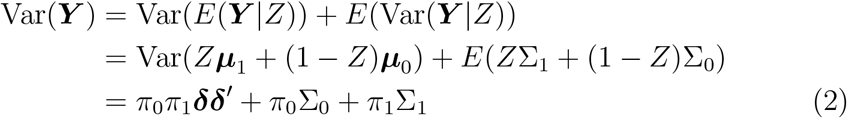

where *Z* is the population indicator variable (*Z* = 0,1) and ***δ*** is ***μ***_1_ − ***μ***_0_ and Σ_*k*_ is the covariance matrix of population *k* = 0, 1. Both covariance matrices Σ_0_, Σ_1_ are identity matrices so Var(***Y***) = *π*_0_*π*_1_***δδ′*** + *I*. Since for any eigenpair (λ, ***e***) of Var(***Y***),

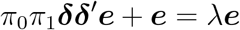

therefore *π*_0_*π*_1_(***δ′e***)***δ*** = (λ − 1)***e***. Thus, the largest eigenvalue is obtained when ***e*** is parallel to ***δ*** and the eigenvalue is

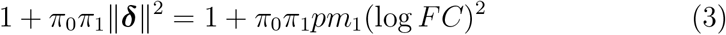

Here the largest eigenvector *PC*1 is ***δ***/||***δ***||. Therefore *PC*1 is the direction between the two population mean vectors ***μ***_1_ and ***μ***_0_.

### *PC*1 in the presence of dropout event

Now consider the dropout mechanism as described in the Method section. Suppose 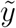 is the gene expression of *y* in the presence of dropout. Then as defined in the Method section, we can write it as

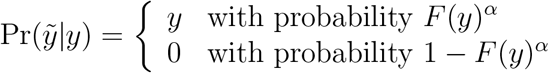

thus, the marginal expectation of 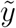 is

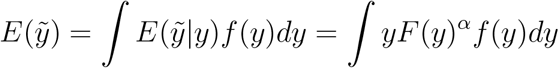

where *f*(*y*) is the density of *y* before dropout event. Since we assume the dropout event is only dependent on the expression value *y*, we can still use the same covariance matrix equation in (2) but with different mean vectors, 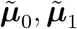 and the covariance matrices, 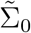 and 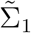. To compute those values, we first need to compute each entry of those vectors. Let 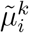 and 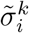 be the mean and the variance of *i*th gene in population *k* then we have

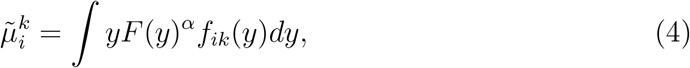

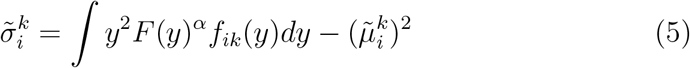

where *f_ik_* is the density of the *i*th gene in Π_*k*_ as in (1). *F* and *α* are same as described in the Method section. From (2), we have the covariance matrix in the presence of dropout as

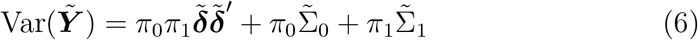

where the *i*th entry of 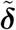 is 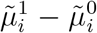 and the *i*th diagonal entry of diagonal matrix 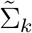 is 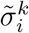 for *k* = 0, 1. Note that 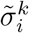 depends on *μ_i_* so 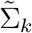 is not necessarily identity matrix.

The main concern is when the largest eigenvector (*PC*1) of (6) is able to separate the two populations of interest. In other words, we need to investigate when *PC*1 **of** (6) **is not orthogonal to** 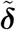, **the population separating direction**.

Now consider a decomposition of 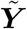 into non-marker and marker genes, say 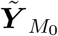 and 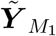. Then the corresponding mean difference vector 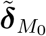 of non-marker set *M*_0_ is zero vector and

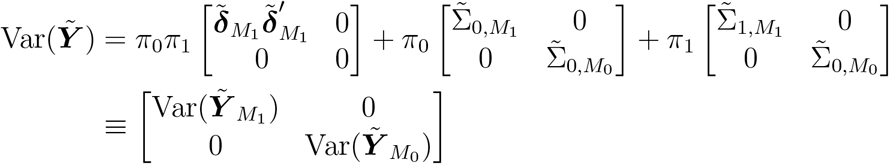

Since Var(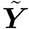) has a block diagonal structure, the eigenvectors of Var(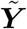) can be obtained separately from Var(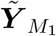) and Var(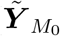) and the two sets of eigenvectors are orthogonal to each other. Therefore, if we let λ_*M*_0__ and λ_*M*_1__ be the largest eigenvalues of Var(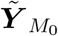) and Var(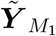), then a necessary condition for *PC*1 separating the two populations (or not orthogonal to 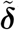) is λ_*M*_1__ > λ_*M*_0__.

We claim that λ_*M*_1__ > λ_*M*_0__ is also a sufficient condition for *PC*1 separating the two populations.

#### Proof.

For simplicity, let 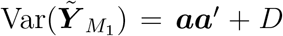 where 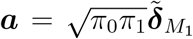 and 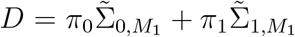 and let ***e*** is the largest eigenvector. Suppose thaty ***e*** and a are orthogonal. Then since ***e*** is an eigenvector,

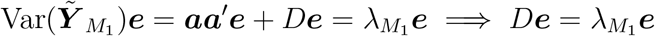

implies ***e*** is an eigenvector of *D*. Since *D* is a diagonal matrix, all eigenvectors are the Euclidean coordinate vectors of *D*, which in turn ***e*** is one of them, say ith coordinate vector. Then the orthogonality assumption implies *i*th entry of ***a, a′ e*** is zero. However it contradicts to the fact that 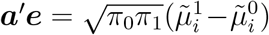 is not zero by construction in (4).

We then investigate the factors that determine λ_*M*_0__ and λ_*M*_1__. Since λ_*M*_0__ is the maximun entry of diagonals 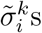 for *i* ∈ *M*_0_ (see (5)), we can see that λ_*M*_0__ is largely affected by the variation of the mean values of individual non-marker genes. For λ_*M*_1__, the characteristic polynomial of Var(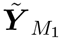) is

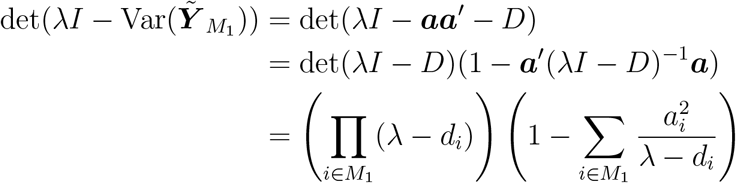

where 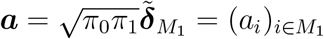 and 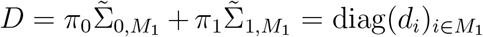. As far as we know, there is no analytic solution for the largest root of the characterstic polynomial. So as a special case we assume *d_i_* = *d* for all *i*. Then for an eigenvalue λ it satisfies

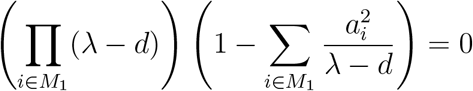

and it implies

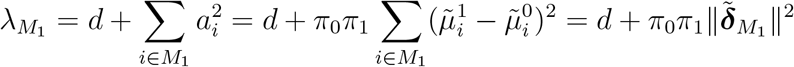

so the magnitude of λ_*M*_1__ is largely determined by 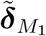 and *π*_1_ (Compare this with (3)).

### Rank of λ_*M*_1__

When λ_*M*_1__ < λ_*M*_0__, the *PC*1 is not informative in separating the two populations of interest as it is orthogonal to 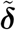. In this case, to detect the population separating direction, we need more than one *PC*s and we might need to develop hypothesis testing setup based on the selected multiple *PC*s. It is obvious that the minimal number of *PC*s required in this scenario is the rank of λ_*M*_1__ among the eigenvalues of Var(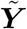).

We calculated the rank of λ_*M*_1__ for our simulation study in Supplementary Table 1 (For simulation results, see https://ncells.shinyapps.io/ncells) where *μ* = 1 and *σ* = 1 of mean gene expression were used. In most conditions, λ_*M*_1__ is larger than λ_*M*_0__, however when *FC* or marker gene fraction *m*_1_ is small, the ranks of λ_*M*_1__ become larger than λ_*M*_0__. In those cases, *PC*1 has no information for separating two populations so one can avoid further simulations to calculate powers for the conditions. Function ncomps in the R package *ncells* was used to calculate the rank of λ_*M*_1__.

**Supplementary Table 1:**
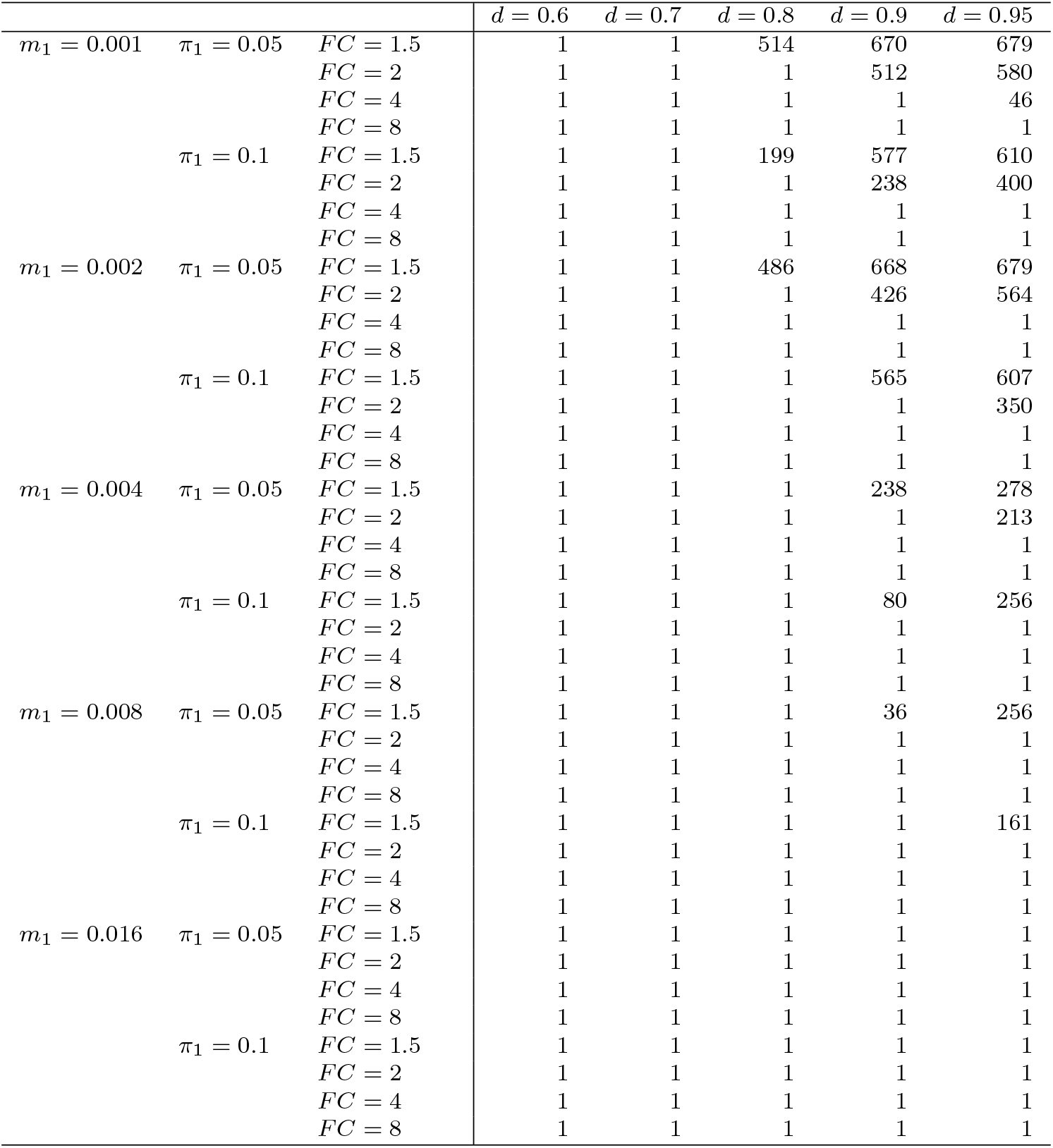
Rank of λ_*M*_1__ among the eigenvalues of Var(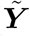). ***μ***_0_ was generated from *N*(0,1) and all entries of ***μ***_1_ are same as ***μ***_0_ except the first 20000 · *m*_1_ entries with difference log *FC*. *m*_1_ is the proportion of marker genes. *π*_1_ is the proportion of minor subpopulation. *FC* is the fold change upregulated in the minor subpopulation. *d* is the overall dropout rate.

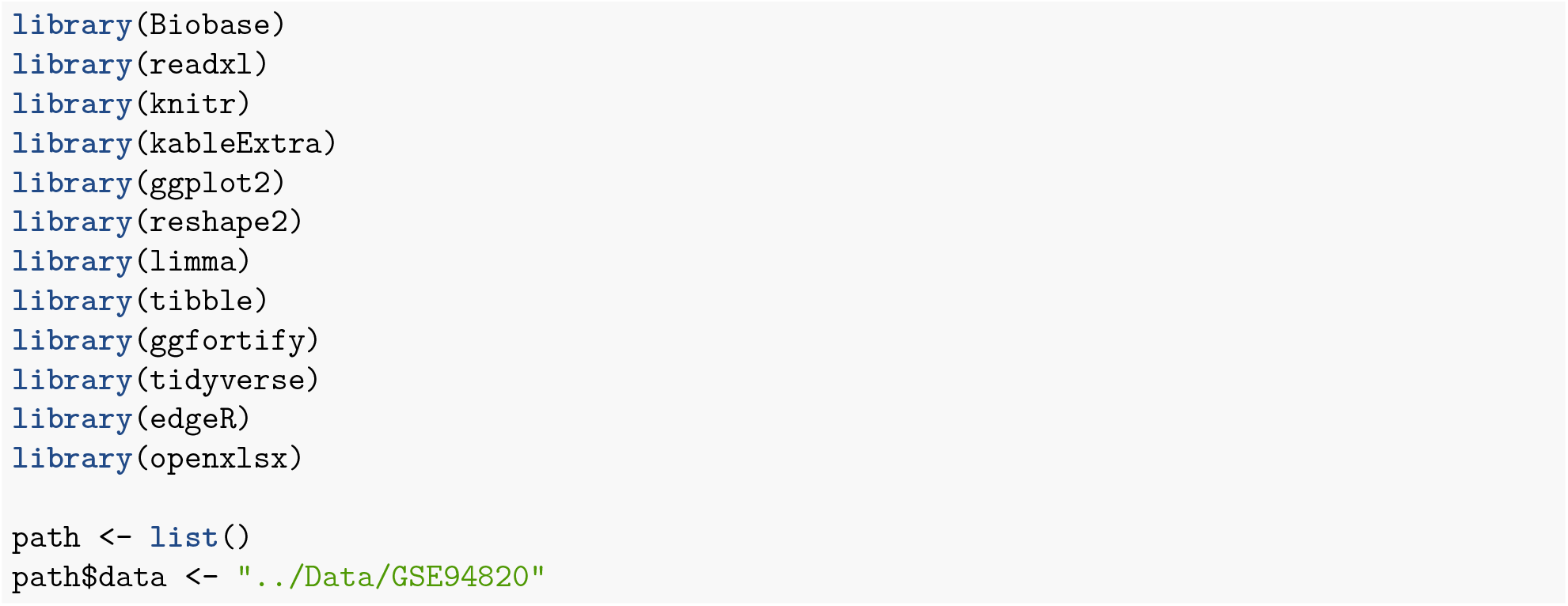

- We downloaded single cell dataset from Gene Expression Omibus (GSE94820). Three known cell populations (two dendritic cell populations, *CD*141 and pDC and one monocyte cell population named as Mono_intermediate) of the dataset were considered for our study. Based on unsupervised clustering, Villani et al. (2017) found 10 clusters from the three populations and three clusters, DC5, Mono3, and Mono4 were selected to investigate as new subpopulations of the three known populations. We first assume those three clusters are new subpopulations and estimate the parameters for simulation using differential expression analysis (edgeR).

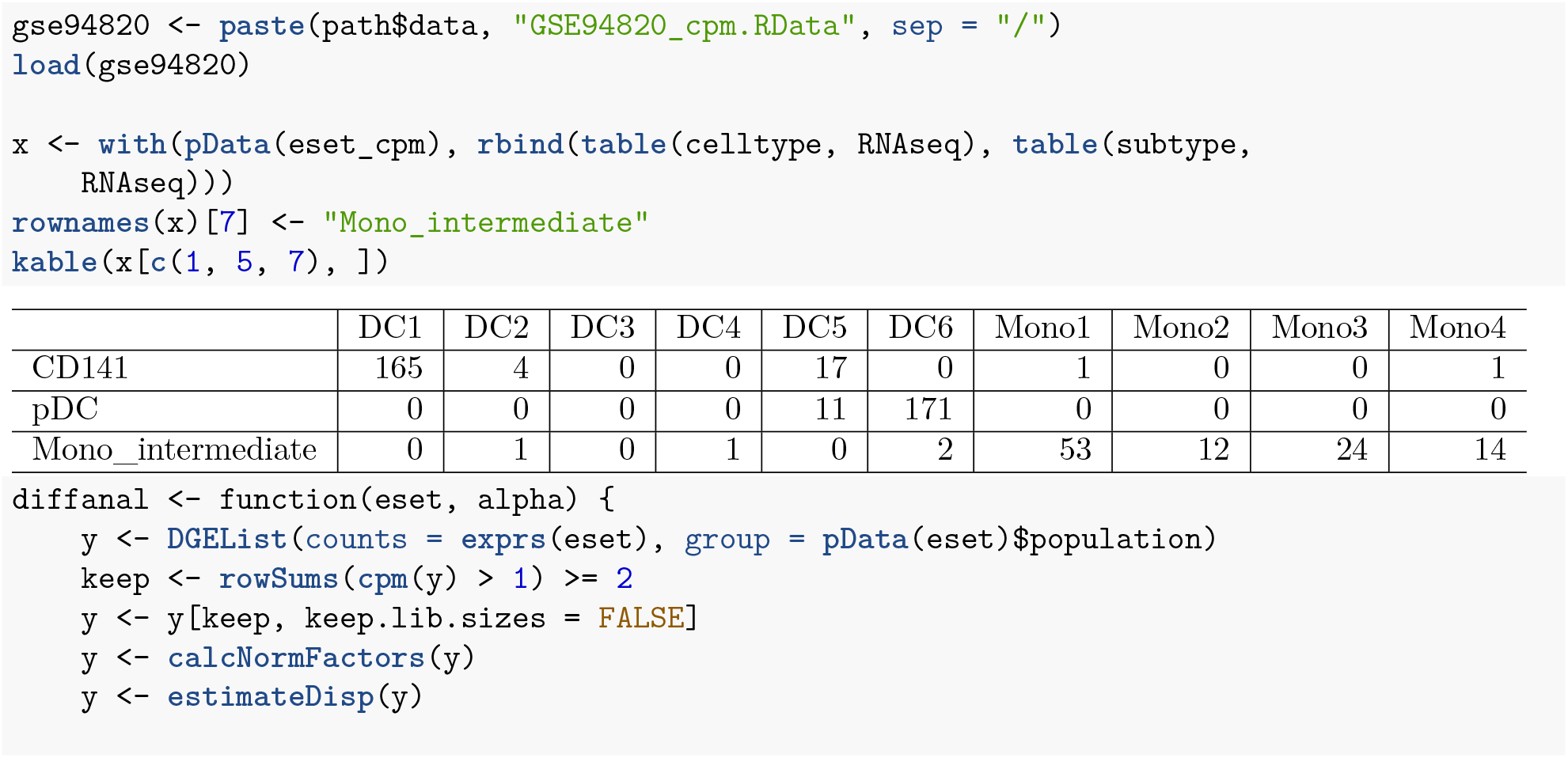
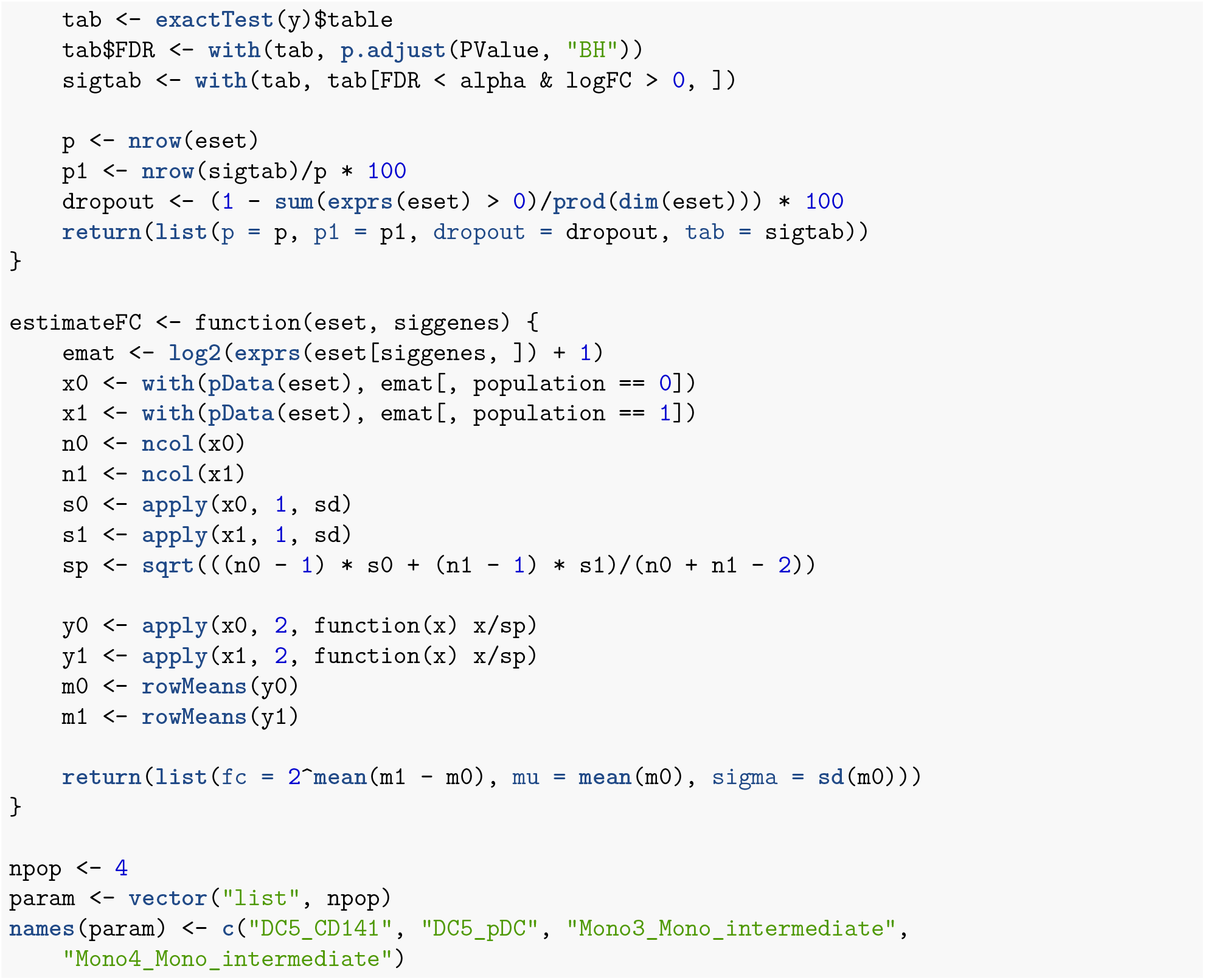

### DC5 subpopulation in CD141 cells

- Total number of cells in CD141: 141
- Number of cells in DC5: 17

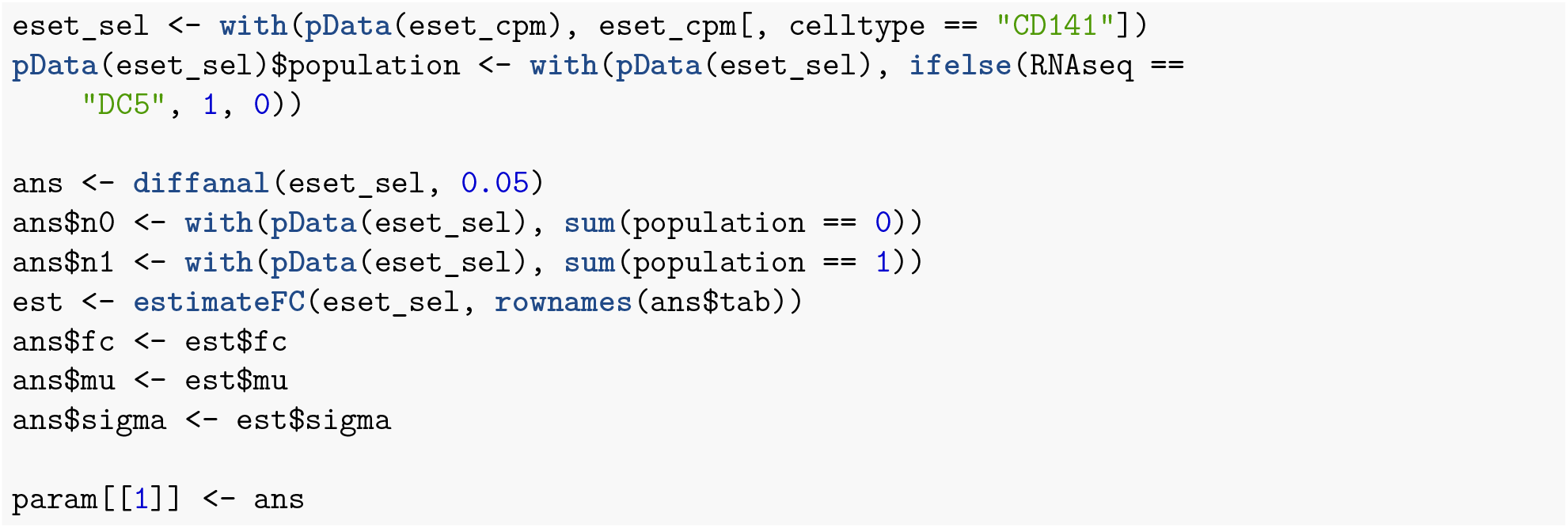

### DC5 subpopulation in pDC cells

- Total number of cells in pDC: 11
- Number of cells in DC5: 182

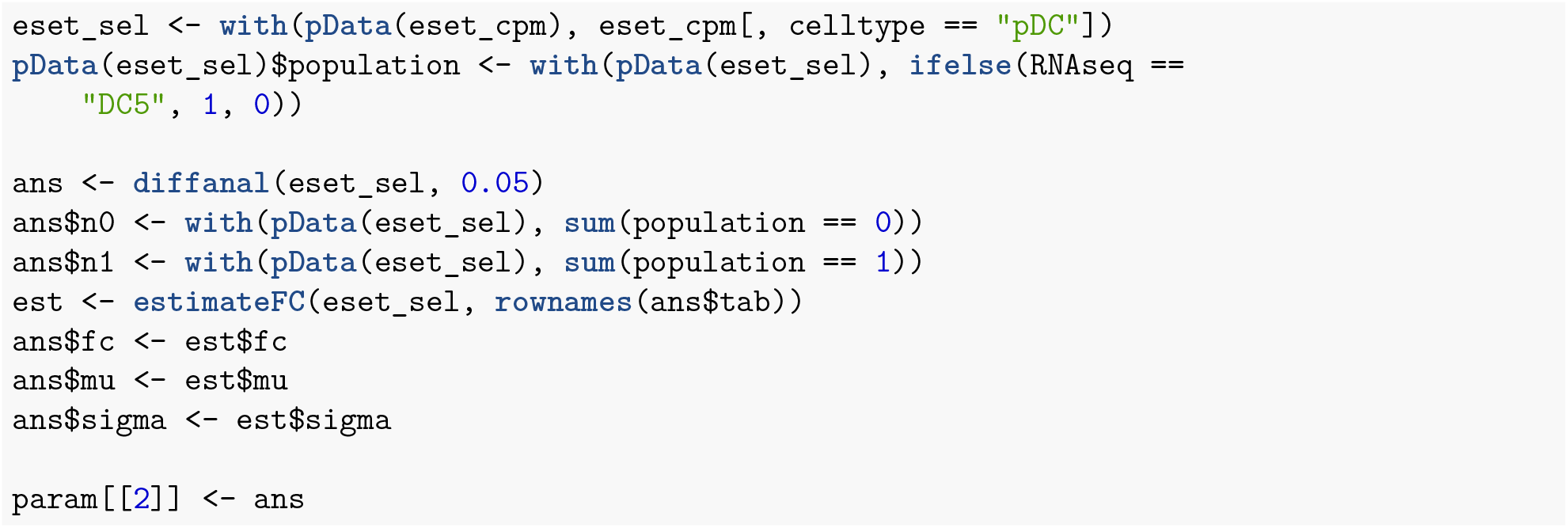

### Mono3 subpopulation in Mono__intermediate cells

- Total number of cells in Mono_intermediate: 107
- Number of cells in Mono3: 24

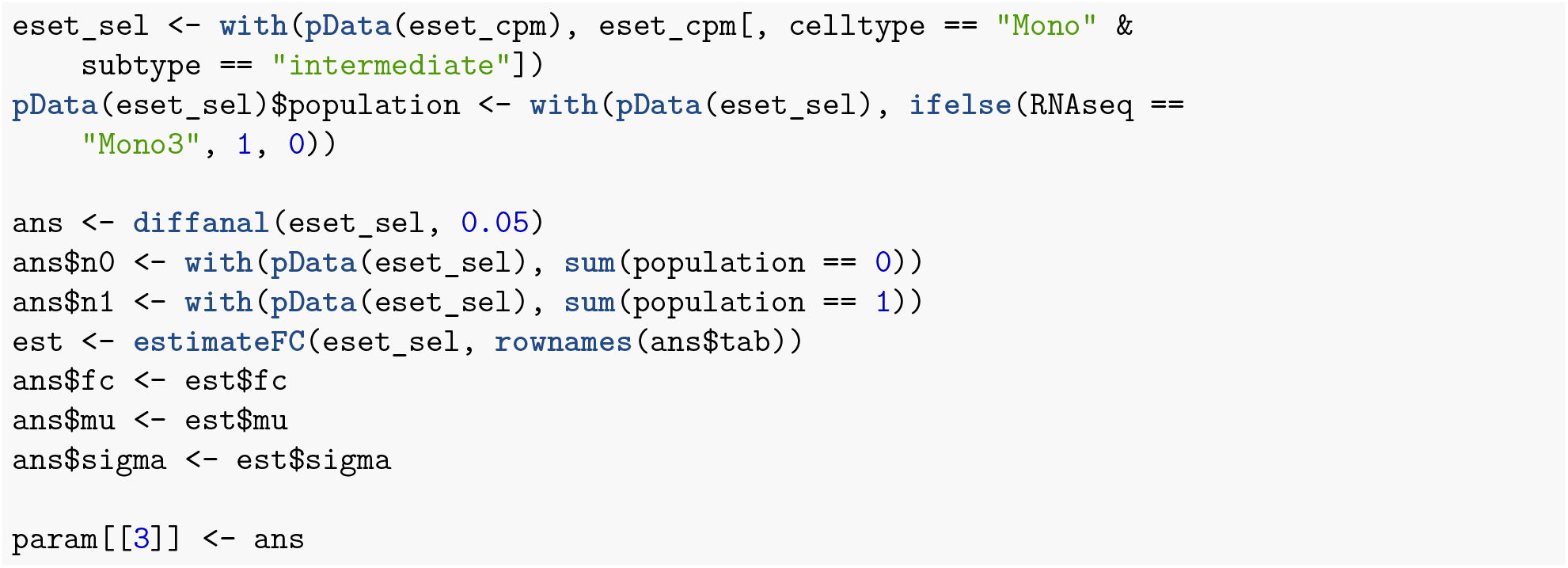

### Mono4 subpopulation in Mono__intermediate cells

- Total number of cells in Mono_intermediate: 107
- Number of cells in Mono4: 14

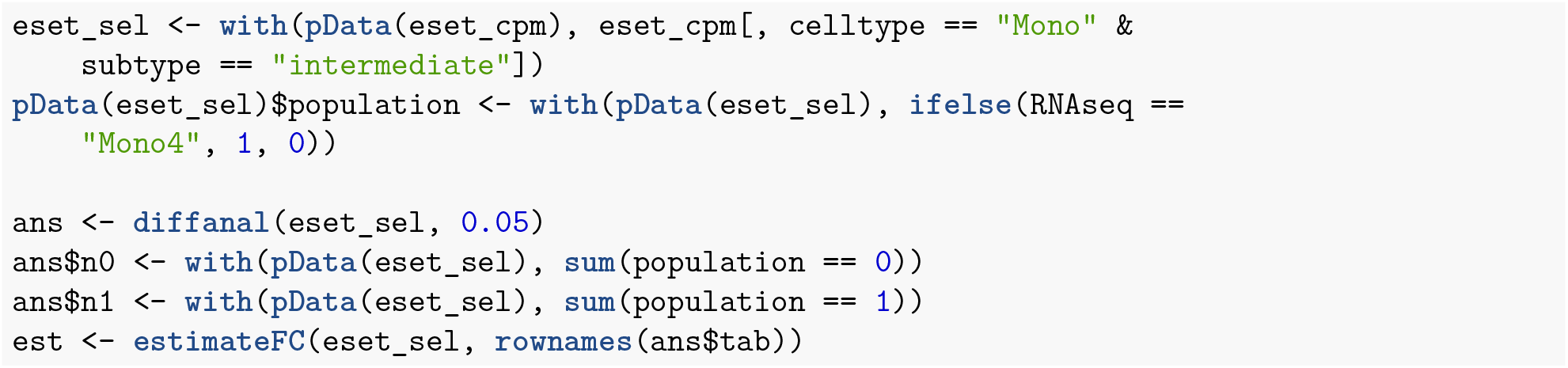
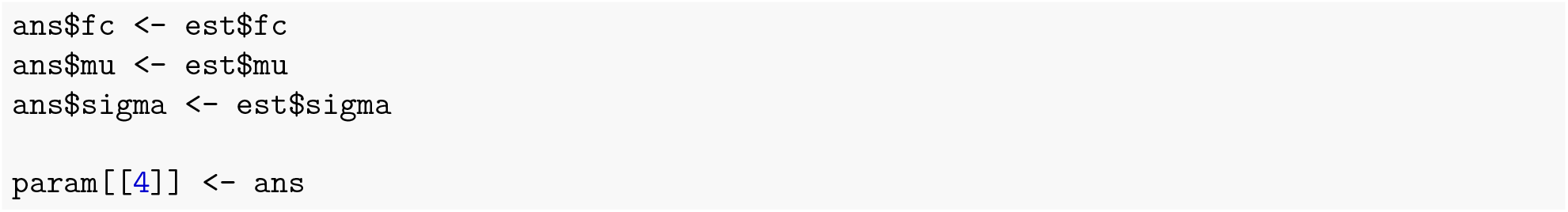

### Summary

- Below is the all the parameters estimated from the single cell dataset.

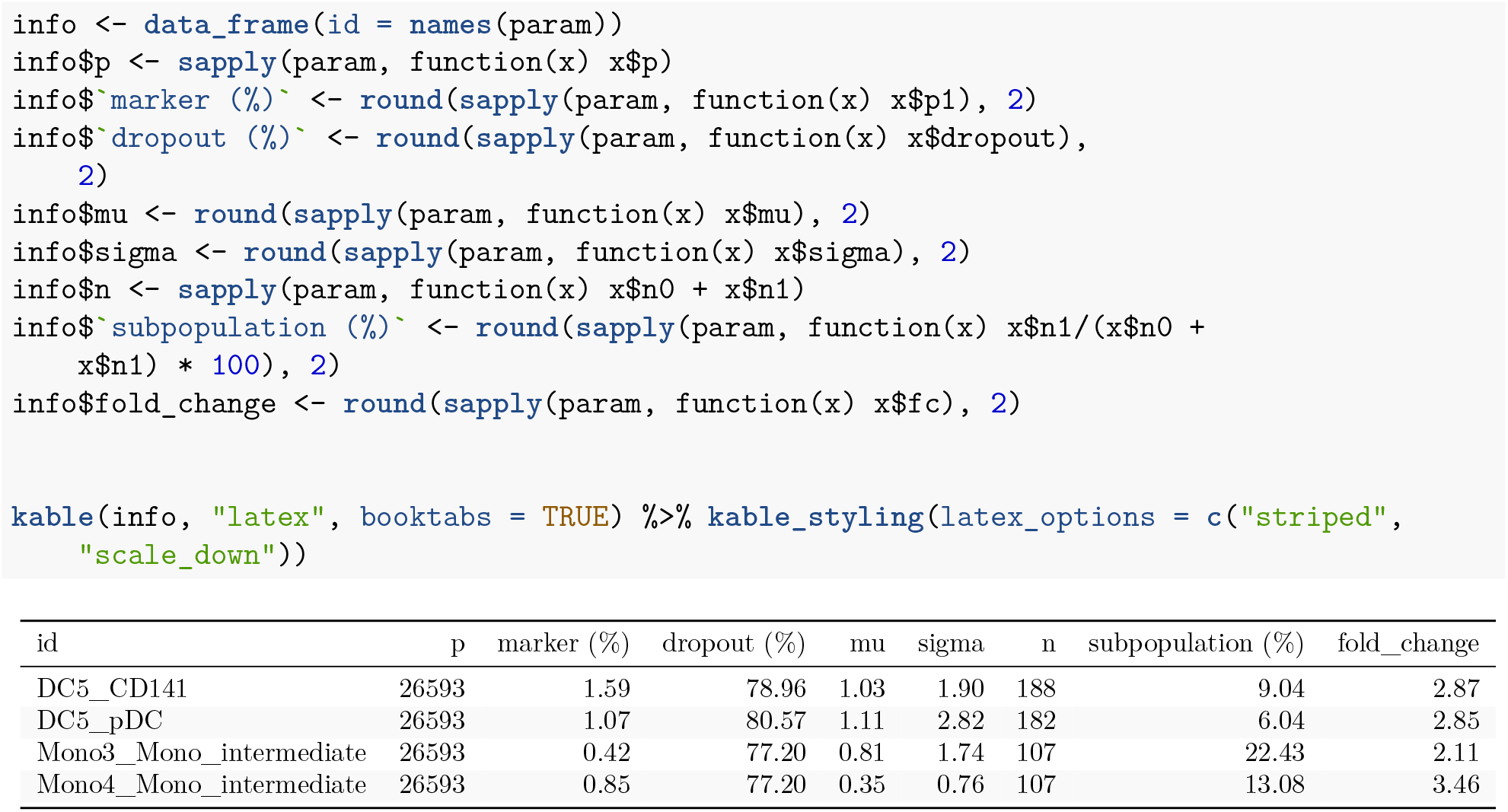

### Sample size plots for four populations

- `res` object in the code below is the output of `ncells` function in `R` package `ncells` version 0.3.0, which was subsequently stored in `RData` file and is called to calculate the power (Current version 0.4.0 directly calculates the power given cut-off values and other conditions). `null`-*population*`.RData` has the output `res` of the function `ncells` with parameter *FC* = 1 and `nonull`-*population*`.RData` has res with parameter *FC* > 1.

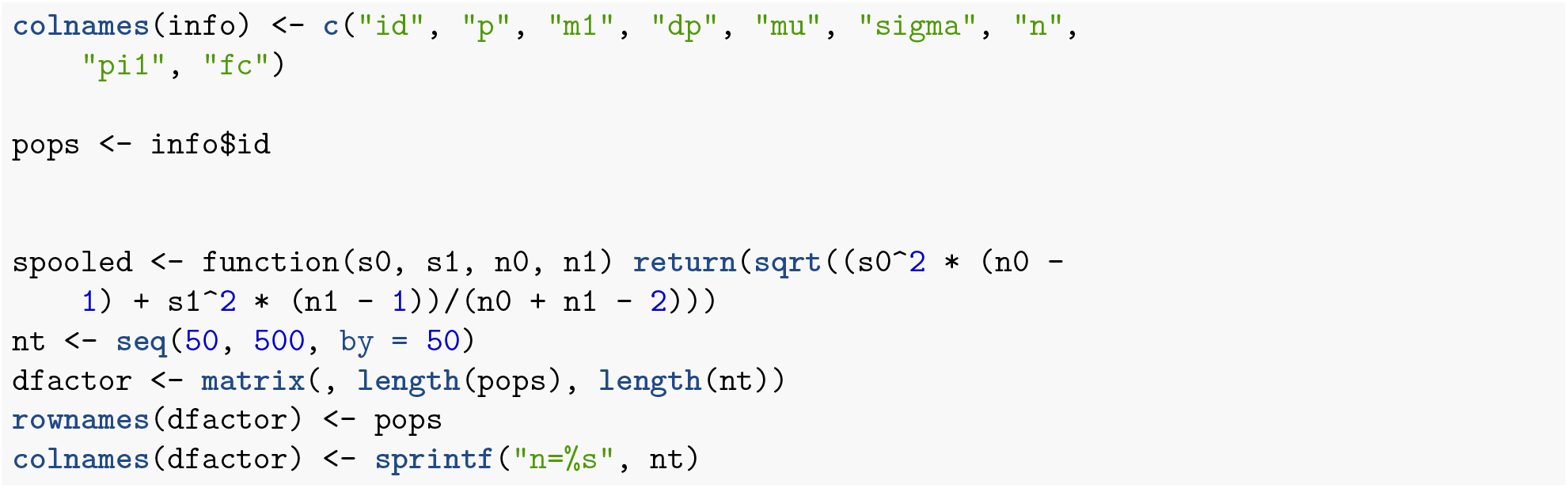
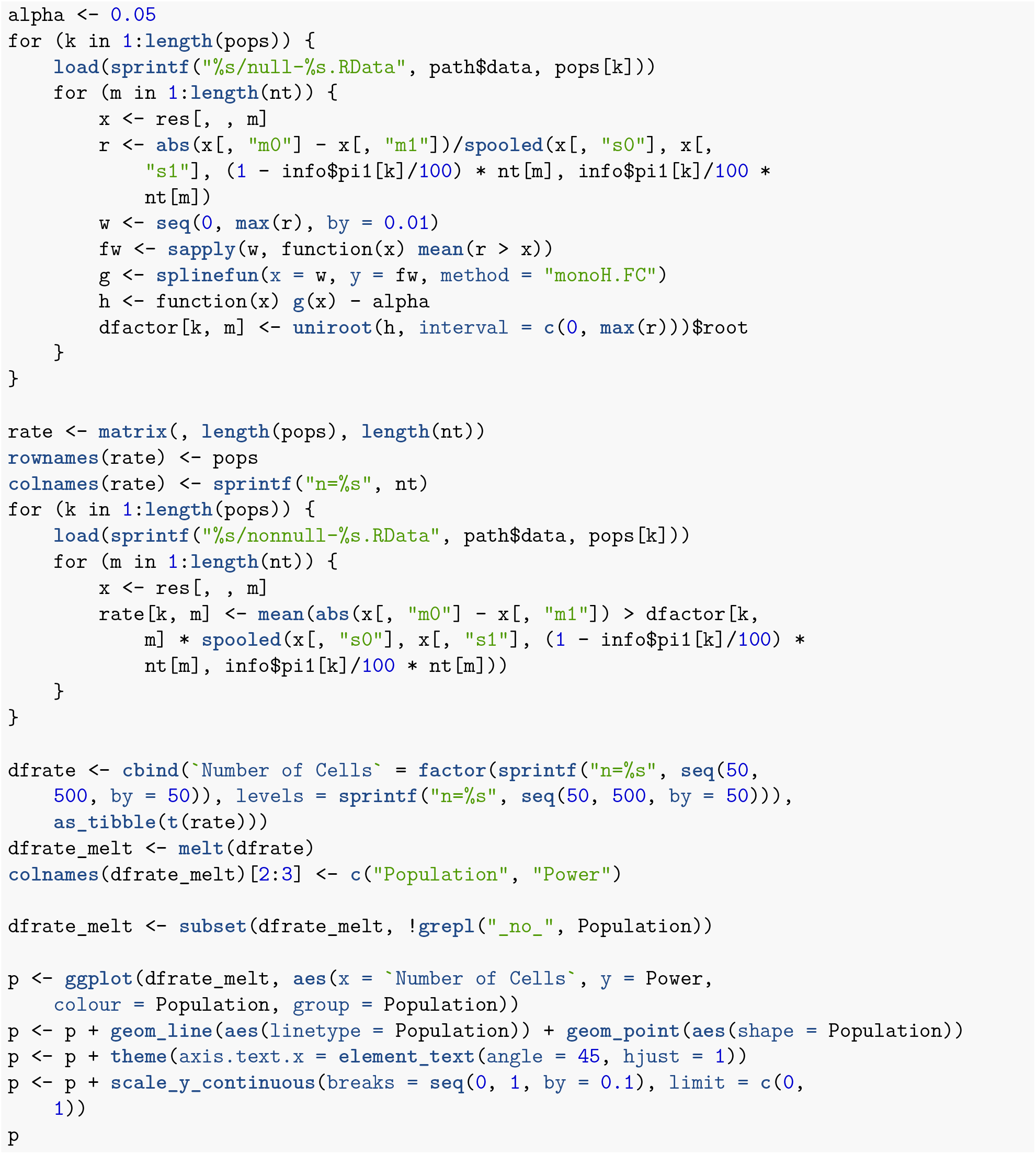

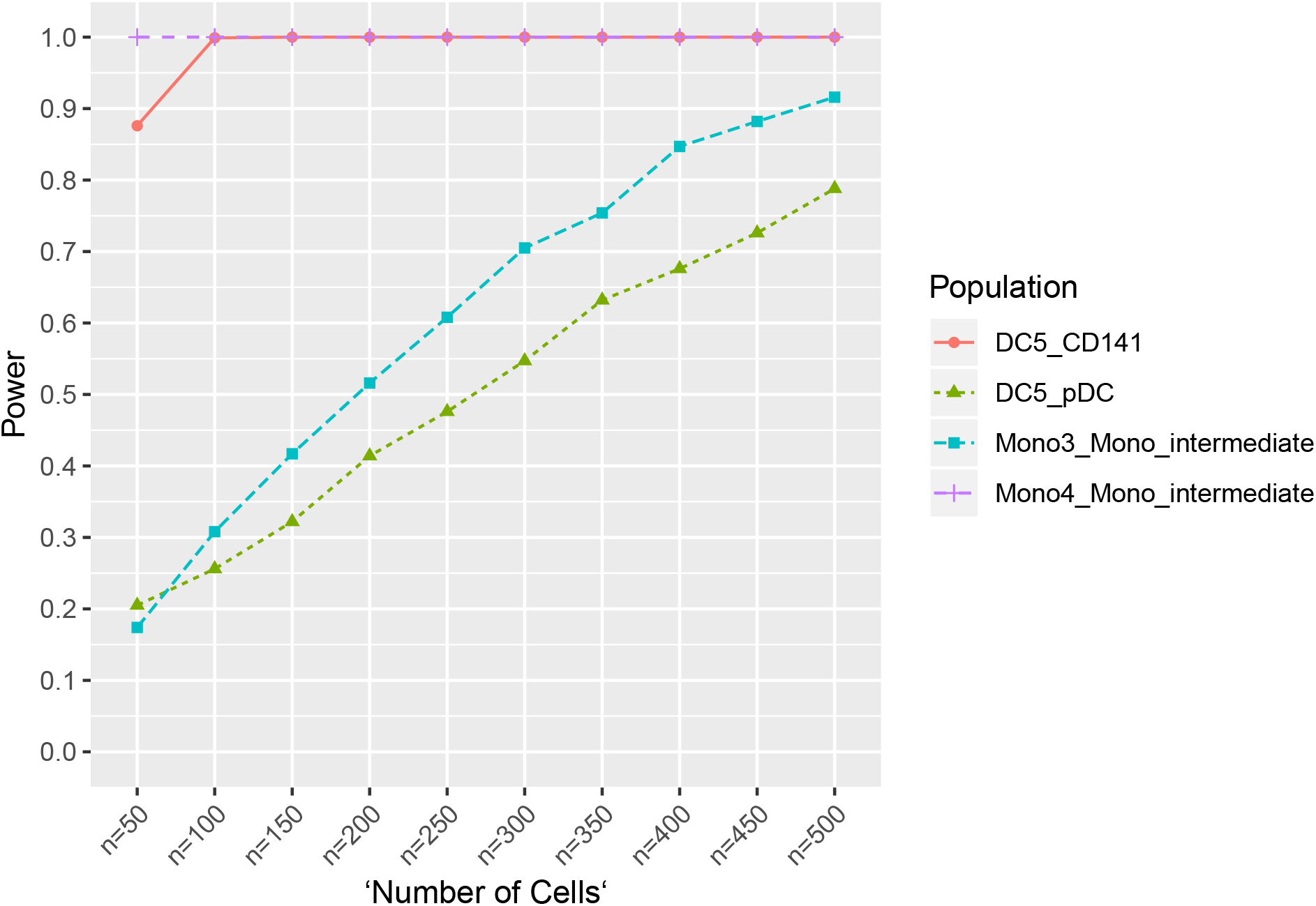

## Notes

https://github.com/TheJacksonLaboratory/ncells

https://ncells.shinyapps.io/ncells

